# Next-Generation Protein Sequencing and individual ion mass spectrometry enable complementary analysis of interleukin-6

**DOI:** 10.1101/2025.02.07.637157

**Authors:** Kenneth A. Skinner, Troy D. Fisher, Andrew Lee, Taojunfeng Su, Eleonora Forte, Aniel Sanchez, Michael A. Caldwell, Neil L. Kelleher

## Abstract

The vast complexity of the proteome currently overwhelms any single analytical technology in capturing the full spectrum of proteoform diversity. In this study, we evaluated the complementarity of two cutting-edge proteomic technologies—single-molecule protein sequencing and individual ion mass spectrometry—for analyzing recombinant human IL-6 (rhIL-6) at the amino acid, peptide, and intact proteoform levels. For single-molecule protein sequencing, we employ the recently released Platinum^®^ instrument. Next-Generation Protein Sequencing^™^ (NGPS^™^) on Platinum utilizes cycles of N-terminal amino acid recognizer binding and aminopeptidase cleavage to enable parallelized sequencing of single peptide molecules. We found that NGPS produces single amino acid coverage of multiple key regions of IL-6, including two peptides within helices A and C which harbor residues that reportedly impact IL-6 function. For top-down proteoform evaluation, we use individual ion mass spectrometry (I^2^MS), a highly parallelized orbitrap-based charge detection MS platform. Single ion detection of gas-phase fragmentation products (I^2^MS^2^) gives significant sequence coverage in key regions in IL-6, including two regions within helices B and D that are involved in IL-6 signaling. Together, these complementary technologies deliver a combined 52% sequence coverage, offering a more complete view of IL-6 structural and functional diversity than either technology alone. This study highlights the synergy of complementary protein detection methods to more comprehensively cover protein segments relevant to biological interactions.

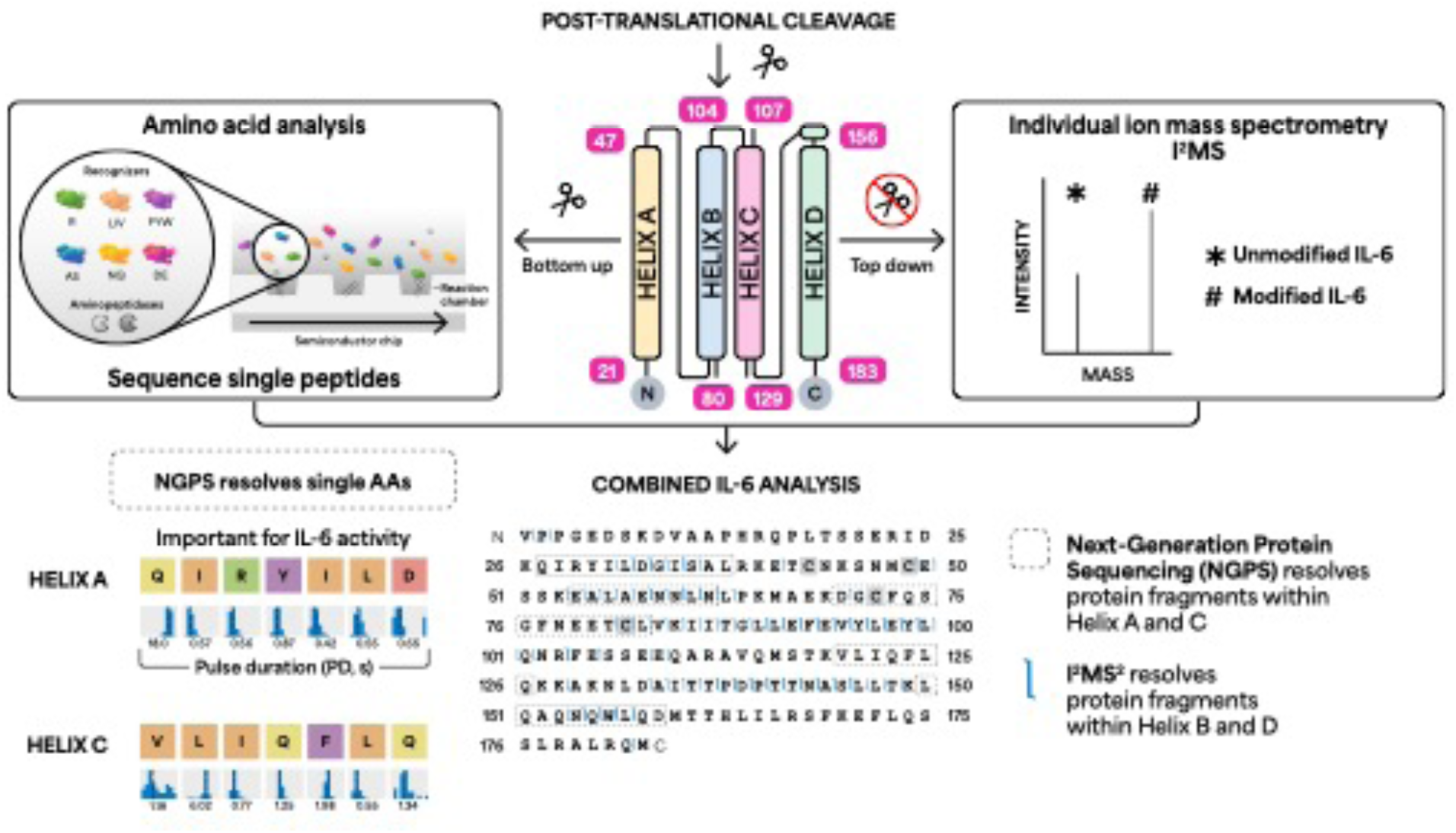

## Introduction

Proteins are fundamental to nearly all biological processes, serving as key cellular components and prime drug targets [1, 2]. Protein-based therapies are among the top-selling pharmaceutical products [3], and recombinant cytokines and biologic inhibitors of cytokine-mediated signaling are used clinically [4, 5]. The structure and function of proteins are determined by their primary amino acid sequence and post-translational modifications (PTMs), making the analysis of these features crucial for drug development and biopharmaceuticals [6, 7].

Rational design and directed evolution are frequently employed to engineer proteins with desirable properties [8, 9]. However, amino acid substitutions, errors in protein synthesis, and variations in expression systems can lead to subtle differences in protein structure and function, necessitating highly sensitive methods for detection [10, 11]. Furthermore, proteins often exist in multiple forms, or proteoforms [12], generated not only by genetic variation but also by alternative splicing and PTMs. Current proteomic technologies struggle to fully capture the complexity and heterogeneity of proteoforms, underscoring the need for innovative solutions [1, 13–16].

Several new technologies offer promising approaches to overcoming these challenges. Next-Generation Protein Sequencing (NGPS) uses engineered protein-based recognizers to sequence peptides with single amino acid resolution, while individual ion mass spectrometry (I^2^MS) is a mass spectrometry technique that measures the intact mass of proteins. Both NGPS and I^2^MS are highly parallelized, single-molecule approaches to analyze protein sequences and PTMs.

Proof-of-concept studies support the enormous potential of single-molecule protein sequencing and analysis [17–19]. The first of these technologies to be commercialized is NGPS on the Platinum and Platinum Pro instruments, which employ fluorophore-labeled N-terminal amino acid (NAA) recognizers that reversibly bind cognate NAAs [20]. These sensitive biophysical interactions produce kinetic signatures that summarize the average sequencing behavior of an ensemble of single peptide molecules, including their primary sequence and PTMs.

The application of individual ion technology to top-down mass spectrometry (TD-MS) provides a >500x improvement in analytical sensitivity, >10x increase in mass range, and >10x higher resolution for the characterization of proteoforms than traditional TD-MS approaches [21, 22]. I^2^MS uses multiplexed orbitrap-based charge detection mass spectrometry (CDMS) to directly measure the charge of individual proteoform ions, producing significantly simplified mass-domain spectra for mixtures of proteoforms without the challenges of deconvolution [23]. Mass differences between the measured protein molecular weight and that of its DNA-predicted sequence may reveal sequence variations and/or PTMs [24]. Further, tandem-MS with the detection of individual ions (I^2^MS^2^) resolves fragment ions from the measured molecular ion to provide sequence analysis across the whole protein, confident protein identification and localization of PTMs [21].

NGPS and I^2^MS utilize distinct fluorescence- or mass spectrometry-based techniques to characterize proteins and their proteoforms. In the context of biopharmaceutical quality control, there is an increasing demand for orthogonal methods—those that employ different physical principles to measure the same property—and for complementary approaches that provide a more comprehensive assessment of protein characteristics [25]. Both I^2^MS and NGPS represent promising single-molecule technologies that meet these needs. Moreover, when used together, NGPS and I^2^MS can provide complementary insights, covering overlapping and distinct protein regions, thus enhancing overall sequence coverage and depth.

Cytokines, such as interleukin-6 (IL-6), play a vital role in immune regulation and are widely used in therapeutic applications [5, 9, 26–28]. The functional versatility and adjustable properties of cytokines have driven the development of engineered variants with optimized pharmacological profiles and enhanced biological activity [9, 29–32]. Beyond genetic modifications, cytokines can undergo alternative splicing [33], post-translational proteolysis [34], and various PTMs [35], all of which can affect their stability and interactions with biological targets [36, 37]. These variations may also differ depending on the expression system used [38, 39]. Understanding and controlling these factors is essential for refining recombinant IL-6 formulations and optimizing their clinical efficacy.

In this study, we use two independent technologies to analyze a recombinant human IL-6 (rhIL-6) sample: NGPS for single-molecule protein sequencing [20] and I^2^MS^2^ for intact mass profiling and readout of top-down fragmentation spectra [22]. By integrating these orthogonal methods for the first time, we show how they complement each other to provide more comprehensive coverage of key regions crucial for IL-6 function and therapeutic potential.

## Materials and Methods

### Peptide sequencing on Platinum

Experiments were conducted in accordance with Library Preparation Kit -Lys-C Data Sheet and Platinum Instrument and Sequencing Kit V3 Data Sheet (February 27, 2024).

Platinum uses bottom-up processing to sequence single peptides and identify proteins. Briefly, the disulfide bonds of rhIL-6 sourced from AcroBiosystems (IL6-H4218) were reduced, enabling alkylation of cysteine residues. Digestion of rhIL-6 with Lys-C endoproteinase [40] generated peptides with C-terminal lysine residues. These peptides underwent diazotransfer [41] and bioconjugation to a macromolecular linker [42, 43]. Conjugated peptides were immobilized in nanoscale reaction chambers on a semiconductor chip for exposure to a mixture of freely diffusing NAA recognizers and aminopeptidases [20]. The sequencing mixture consisted of six NAA recognizers that target 13 NAAs. During on-chip sequencing, fluorophore-labeled NAA recognizers reversibly bind cognate NAAs, producing recognition segments (RSs) and fluorescence properties that are captured by the semiconductor chip. To carry out the sequencing process, aminopeptidases cleave the peptide bond and expose the subsequent NAA for recognition.

### Analysis of sequencing data; analysis versions

Primary Analysis v2.5.1, Peptide Alignment v2.3.0, and Protein Inference v2.5.2 were used for cloud-based analysis of sequencing data. Details can be found in the <u>Platinum Analysis</u> <u>Software Data Sheet</u> (February 2, 2024). The Primary Analysis workflow is the first step in processing data which characterizes the apertures across the chip based on peptide loading, recognizer activity, recognizer reads, and recognizer read lengths.

### Peptide Alignment v2.3.0 workflow

For the Peptide Alignment workflow, a reference sequence is required to call amino acids from the recognizer reads. Reads from the sequencing data were aligned to the FASTA reference of mature rhIL-6. Peptides were aligned based on the correspondence of observed recognition segments to the expected reference profile, using recognizer identity.

Peptide Alignment workflow also computes a false discovery rate (FDR) for each aligned peptide. This calculation is adapted from methods used in peptide identification by mass spectrometry [44], based on decoy peptide matching. Thus, FDR represents the relative number of alignments to the reference peptide sequence versus the total number of off-target alignments, such as scrambled sequences with the same length as the target peptide.

### Protein Inference workflow v2.5.2

A pre-defined reference set of 8,076 human proteins was used to infer proteins from unknown samples and confirm the identity of samples. The proteins in this reference panel span 10–70 kDa and contain at least three in silico LysC-digested peptides with three unique, visible residues. Inferred proteins are ranked by their respective Inference Score. The Inference Score is a natural log calculation of the FDR associated with the inferred protein.

### Sample preparation and data collection for I^2^MS and I^2^MS^2^ analysis

RhIL-6 (AcroBiosystems, IL6-H4218) was reconstituted in Optima^TM^ LC/MS grade water (Fisher Scientific, W64), aliquoted, and stored at -80°C until used following the manufacturer’s recommendations. Samples were analyzed with the SampleStream Platform (Integrated Protein Technologies, Evanston, IL) [45] coupled to a modified Q-Exactive HF (Thermo Fisher Scientific; Bremen, Germany) mass spectrometer. Briefly, rhIL-6 was diluted to ∼ 0.8 µM with water and transferred to a low-retention autosampler vial (Waters, 186009186). For each injection,10 µL was buffer exchanged with SampleStream into 80 µL denaturing buffer (70:30 water:acetonitrile with 0.2% formic acid), deposited into a clean vial to mix, and then 75 µL aspirated to infuse at ∼ 0.1 µM. SampleStream parameters include a 125 µL push volume, 125 µL focus volume, 225 µL focus flow rate, 60°C flow cell temperature, and a 5 kDa molecular weight cut-off membrane. Source conditions include a custom nano-electrospray emitter (CoAnn Tech, TIP36007540-10), 1.8 kV spray voltage, 1.0 µL/min flow rate, and 320°C inlet capillary temperature. Instrument parameters include RF 50%, 5e6 AGC target, 120,000 resolution, 1 µscan, -1 kV central electrode voltage, 0.3 (arb) trapping gas pressure setting, 600-2500 *m/z* scan range, and a 78-minute acquisition length (4583 scans). Injection time was determined via automated ion control (AIC) to remain on the individual ion level [46].

For I^2^MS^2^ experiments, a charge state for each precursor was selected using The Fisher, a tool developed internally by the Kelleher group [47]. Briefly, ions within a 0.8 *m/z* isolation window width and 20 Da mass window width centered on a precursor mass were counted as ions corresponding to either the desired precursor or other species. The charge state which maximized the number of desired precursor ions and minimized the number of ions from other species was selected for isolation. Precursors were isolated and fragmented via higher-energy collisional dissociation (HCD) normalized to charge state. Fragment ions were measured within a 150-2500 *m/z* scan range. Normalized collisional energy (NCE) and injection time were set manually to optimize fragmentation and ion counts.

### I^2^MS and I^2^MS^2^ data processing and analysis

The I^2^MS charge assignment and ion mass determination process has been previously described [22, 48]. The *m/z* of individual ions were determined through normal Fourier-transform Mass Spectrometry (FTMS) analysis, while the charge of the ion was determined using the summation of its ion signal as a function of time (STORI plot) [48]. Once the quantized charge and *m/z* values of the ion were assigned, the mass of the species was calculated and utilized to create a true mass spectrum.

A fragment search against the precursor amino acid sequence and expected PTMs was carried out using TDValidator (Proteinaceous, Inc., Evanston, IL). Fragment ions were identified by matching their isotopic distributions to theoretical isotopic distributions generated using an averagine model [49] and the Mercury7 [50] algorithm. To make ions searchable in TDValidator, neutral mass I^2^MS spectra were transformed into theoretical +1 (M+H) distributions. All fragment ions were identified within a ±10 ppm tolerance of their theoretical values for the isotopic distribution (Max PPM tolerance) and ±5 ppm tolerance for isotopologues within the same distribution (Sub PPM tolerance). Other search metrics included a +1 charge state, 1.5 S/N cutoff, and 0.01 score cutoff. Spectra were manually curated to remove poor fragment ion matches. To calculate P-scores, fragment monoisotopic masses were generated with the THRASH algorithm in TDValidator using a 1.5 S/N cutoff, charge +1, 30,000 Da maximum mass, and 0.9 minimum RL value and uploaded to ProSight Lite (http://prosightlite.northwestern.edu/) with 10 ppm tolerances.

## Results and Discussion

To evaluate the ability of each proteomic technology to detect key regions of rhIL-6, we first performed NGPS of rhIL-6 using the Platinum instrument (**Figure 1A**). The Platinum platform includes three components: kits for bottom-up protein processing and sequencing; a benchtop Platinum instrument that accommodates semiconductor chips for sequencing of polypeptides (**Figure 1B and C**); and cloud-based software for analysis of sequencing data (**Figure 1D and E**).

**Figure 1.**
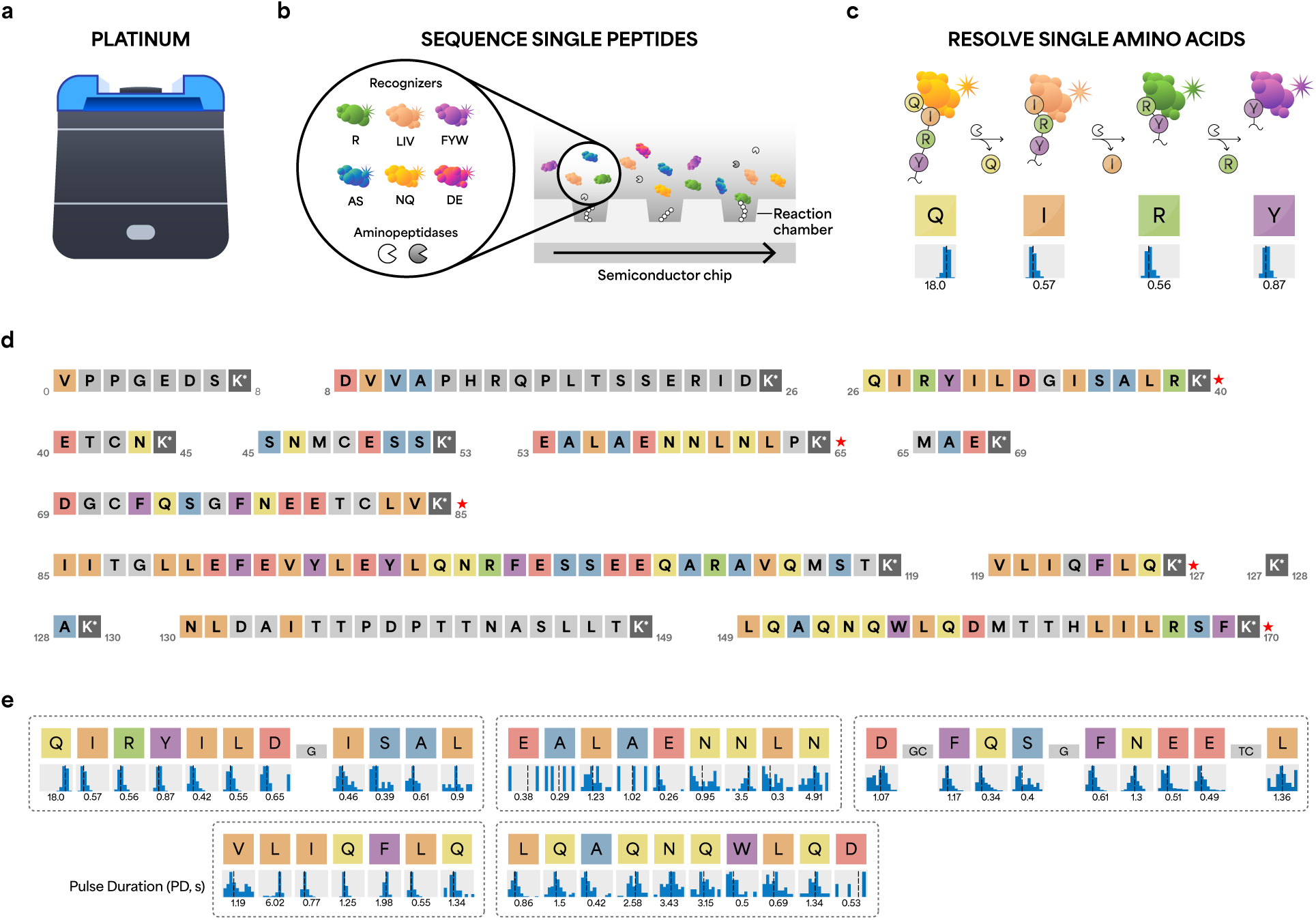
Next-Generation Protein Sequencing (NGPS) of IL-6 with the Platinum instrument. (**a**) The Platinum instrument sequences single peptide molecules with single amino acid resolution. (**b**) Sequencing kits include semiconductor chips, aminopeptidases, and six dye-labeled NAA recognizers that reversibly bind 13 target NAAs. (**c**) Binding of dye-labeled NAA recognizers generates kinetic information indicating which amino acid is being detected. (**d**) Results of *in silico* Lys-C digestion of mature IL-6. Missing C-terminal residues 171-183 lack a lysine residue (K) and thus are not amenable for peptide capture and sequencing on Platinum. Colored boxes indicate potential recognition events. Gray boxes indicate amino acids not amenable to recognition. Red asterisks indicate automated alignments of sequencing data to reference sequence. (**e**) NGPS of rhIL-6 on Platinum detects five IL-6 peptides (dotted rectangles). NAA recognizers elicit kinetic signatures such as pulse duration (PD), which reflects the affinity between specific recognizers and NAAs. PD histograms represents the statistical distribution of kinetic data for all pulses associated with a specific residue and supports inference of the corresponding amino acid. Values are reported as the median of the mean across reaction chambers.

For on-chip sequencing, surface immobilized peptides are exposed to a mixture of freely diffusing NAA recognizers and aminopeptidases (**Figure 1B**) [20]. Six NAA recognizers, labeled with different fluorophores, reversibly bind 13 target NAAs (**Figure 1B**) and elicit characteristic pulsing patterns upon binding to each NAA (**Figure 1C**). The semiconductor chip converts fluorescence signal into digital readouts, enabling real-time sequencing of single peptide molecules in parallel [20]. Cycles of binding and cleavage proceed to sequentially reveal the order of NAAs and enable identification of the peptide sequence [20].

For analysis of sequencing data, only peptides that meet specific thresholds (see **Methods** section) are eligible for high-confidence alignment to the reference rhIL-6 sequence (**red asterisks, Figure 1D**). Based on these criteria, three rhIL-6 peptides (V1PPGEDSK8, E41TCNK45, and M66AEK69) are ineligible for alignment. In addition, the C-terminal segment 171 – 183 (E171FLQSSLRALRQM183) does not contain K and thus is not amenable to conjugation and on-chip sequencing. NGPS sequences five rhIL-6 peptides (**Figure 1E**) and determines the identity of 46/183 single amino acids, indicating ∼25% amino acid level coverage within rhIL-6 (**Figure 1E**).

To support high-confidence identification of IL-6, we also examined the false discovery rate (FDR) for each peptide and used Protein Inference analysis (**see Methods section**) to determine the specificity of IL-6 mapping relative to a protein panel. Platinum analysis outputs FDR for each sequenced peptide using a target decoy approach that is analogous to methodologies employed in MS [44]. FDR scores for each peptide are Q27IRYILDGISALRK40 (FDR: 0.0), V120LIQFLQK127 (FDR: 0.0), D70GCFQSGFNEETCLVK85 (FDR: 0.02), E54ALAENNLNLPK65 (FDR: 0.07), and L150QAQNQWLQDMTTHLILRSFK170 (FDR: 0.09). Interestingly, an FDR score of 0.0 was computed for peptide V120LIQFLQK127, despite contiguous isobaric residues leucine (L) in position 2 and isoleucine (I) in position 3 (**Figure 1E**). The recognizer for branched chain NAAs (LIV) exhibits differential pulse duration (PD) profiles for L (6.02 s) and I (0.77 s), discerning the order of L and I residues with the same mass. This example demonstrates that a single NAA recognizer can differentiate NAAs with similar physiochemical properties on the basis of differences in PD. In addition to FDR, we also used the Protein Inference analysis software (see Methods), which screens the sequencing data against a reference panel of 8,076 proteins. IL-6 was identified as the top protein hit with 99.99% confidence (**Supplementary Figure 1**). These results demonstrate NGPS confidently identifies rhIL-6 with single amino acid resolution.

Next, we deployed the same rhIL-6 sample for intact mass measurement via I^2^MS. I^2^MS accurately identifies unmodified rhIL-6 (∼20.8 kDa) and several higher molecular weight proteoforms up to ∼22 kDa (**Figure 2A**). Human IL-6 has been shown to undergo a number of PTMs including O- and N-linked glycosylation and phosphorylation [35]. O-glycans have been reported in species under 25 kDa, while heavier species up to 29 kDa have been reported to contain N-glycans, though few reports detail the composition or localization of IL-6 O-glycans [51]. Using the intact masses of these proteoforms, the composition of the putative O-glycans was determined by searching the GlyGen database [52] for those glycans corresponding to the observed intact mass shift, and their chemical formula and mass were calculated using the publicly available NIST Glyco Mass Calculator [53] (**Supplementary Table 1**). Thus, I^2^MS enables the observation of intact proteins and proteoforms, which is distinct from the data produced by NGPS.

**Figure 2.**
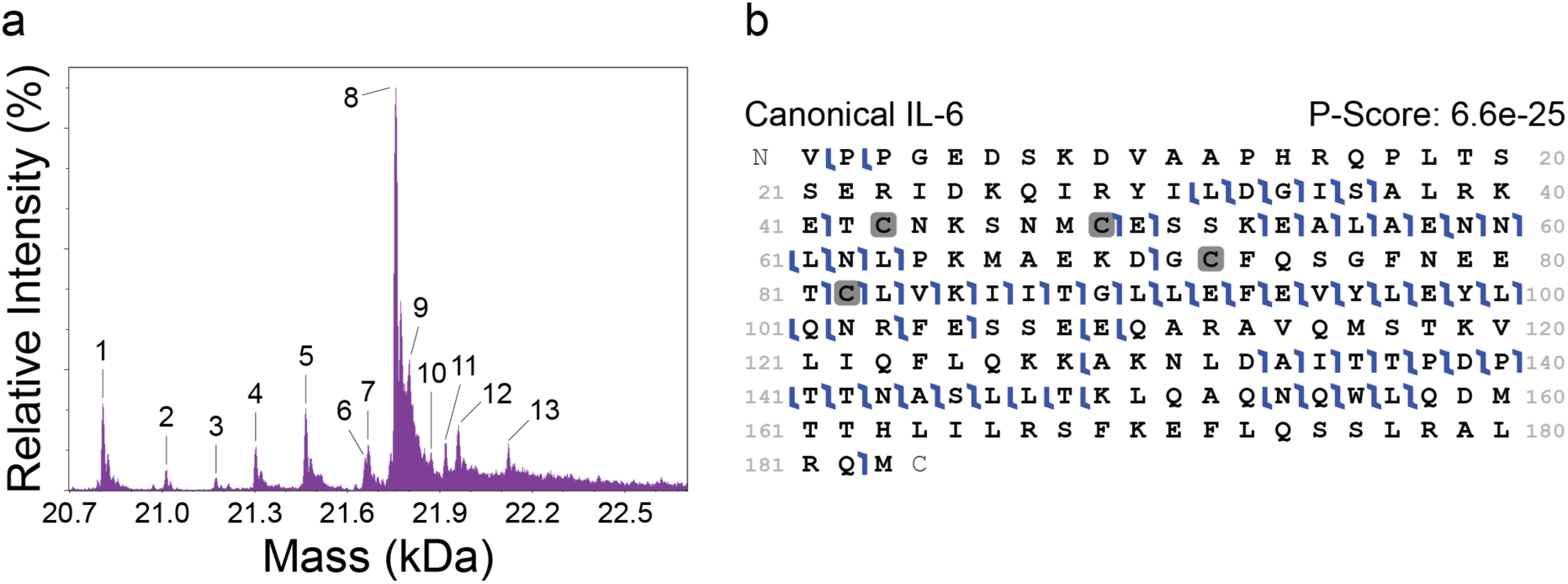
I^2^MS detects intact rhIL-6 proteoforms. I^2^MS^2^ covers broad regions of rhIL-6 primary structure. (**a**) The application of I^2^MS to TD-MS enables measurement of the intact mass of intact protein ions of canonical rhIL-6 (∼20.8 kDa). Higher mass proteoforms indicate the addition of PTMs. (**b**) TD-MS provides broad sequence coverage of canonical rhIL-6 (PFR 1) by higher energy collision dissociation (HCD), achieving 39% sequence coverage.

We then isolated and fragmented discrete rhIL-6 proteoforms via I^2^MS^2^. Using higher energy collision dissociation (HCD), I^2^MS^2^ achieves 39% fragmentation coverage of the unmodified form of rhIL-6, with shared and distinct regions identified by NGPS (**Figures 2B and 1E**). Of the 12 specified higher mass species, 7 more abundant species showed satisfactory ion counts of the desired precursor with minimal co-isolation of other species (PFRs 4, 5, 8, 9, 11, 12, 13) using The Fisher, an in-house tool for determining I^2^MS^2^ isolation windows, on the intact I^2^MS^2^ spectrum. I^2^MS^2^ results of these selected higher mass species are consistent with putative O-glycans with compositions of N-acetyl hexosamine, hexose, deoxyhexose, and sialic acid variably located towards the C-terminus in Helix D and the adjacent C-D loop. Localization of the O-glycans at specific serine or threonine residues along the base rhIL-6 sequence was predicted using the OGP repository [54]. Sites T6, T48, S49, T117, T147, T166, T170, T171, and T177 generated a probability for O-glycosylation greater than 0.5 and thus were used as potential O-glycan locations. Canonical rhIL-6 and select higher mass species with their respective O-glycan composition and localization are supported with fragment map P-Scores ranging from 6.6E^-25^ to 2.3E^-04^ (**Supplementary Figures 2-9**). The O-glycan localization (T147, T166, and T177 across all glycoforms) which generated the smallest P-Score for each proteoform is reported. O-glycosylation located towards the C-terminus of human IL-6 is consistent with that reported in lung adenocarcinoma cells isolated from malignant pleural effusion [51].

Both NGPS and I^2^MS^2^ detect the E54ALAEAENNLNL63 sequence region (**Figures 1E** and **2B**). Also, while NGPS provides single amino acid resolution of the D70GCFQSGFNE79 fragment, only a single residue cleavage was observed within this peptide by I^2^MS^2^ (**Figures 1E, 2B**). Similarly, NGPS reports the sequence of the V120LIQFLQK127 peptide (**Figure 1E**), which also contains no residue cleavages by I^2^MS^2^ (**Figure 2B**).

For the region encompassing Q27IRYILDGISAL38 (**Figures 1E, 2B**), NGPS and I^2^MS^2^ provide complementary coverage. While NGPS enables single amino acid resolution of 11/12 amino acids (**Figure 1E**), I^2^MS^2^ detects fragment ions on either side of the glycine (G) residue, which is not detected by NGPS due to the lack of a G recognizer (**Figure 1B**). While NGPS does not currently provide a C recognizer, I^2^MS^2^ detects C-containing fragments C43NKSNMCESSK53 and C82LVKIITGLLEFEVYLEYLQNR103 (**Figure 2B**).

In addition to this 22 amino acid-long internal fragment, I^2^MS^2^ also provides near complete amino acid coverage across N131LDAIITTPDPTTNASLLTK150 (**Figure 2B**). Both NGPS and I^2^MS^2^ sequence the C-terminal region of rhIL-6 encompassing L150QAQNQW156 (**Figures 1E, 2B**), which contains the only tryptophan (W) in IL-6 [55, 56]. Interestingly, amino acids within this fragment have been implicated in IL-6 binding interactions with receptors.

Overall, our results demonstrate NGPS and TD-MS provide sequence information for overlapping and distinct regions within rhIL-6. To place these sequencing results in the context of IL-6 tertiary structure and function, we mapped the amino acid regions detected by NGPS (**Figure 1E**) and I^2^MS^2^ (**Figure 2B**) using an IL-6 crystal structure as a reference [57] **(Figure 3A).**

**Figure 3.**
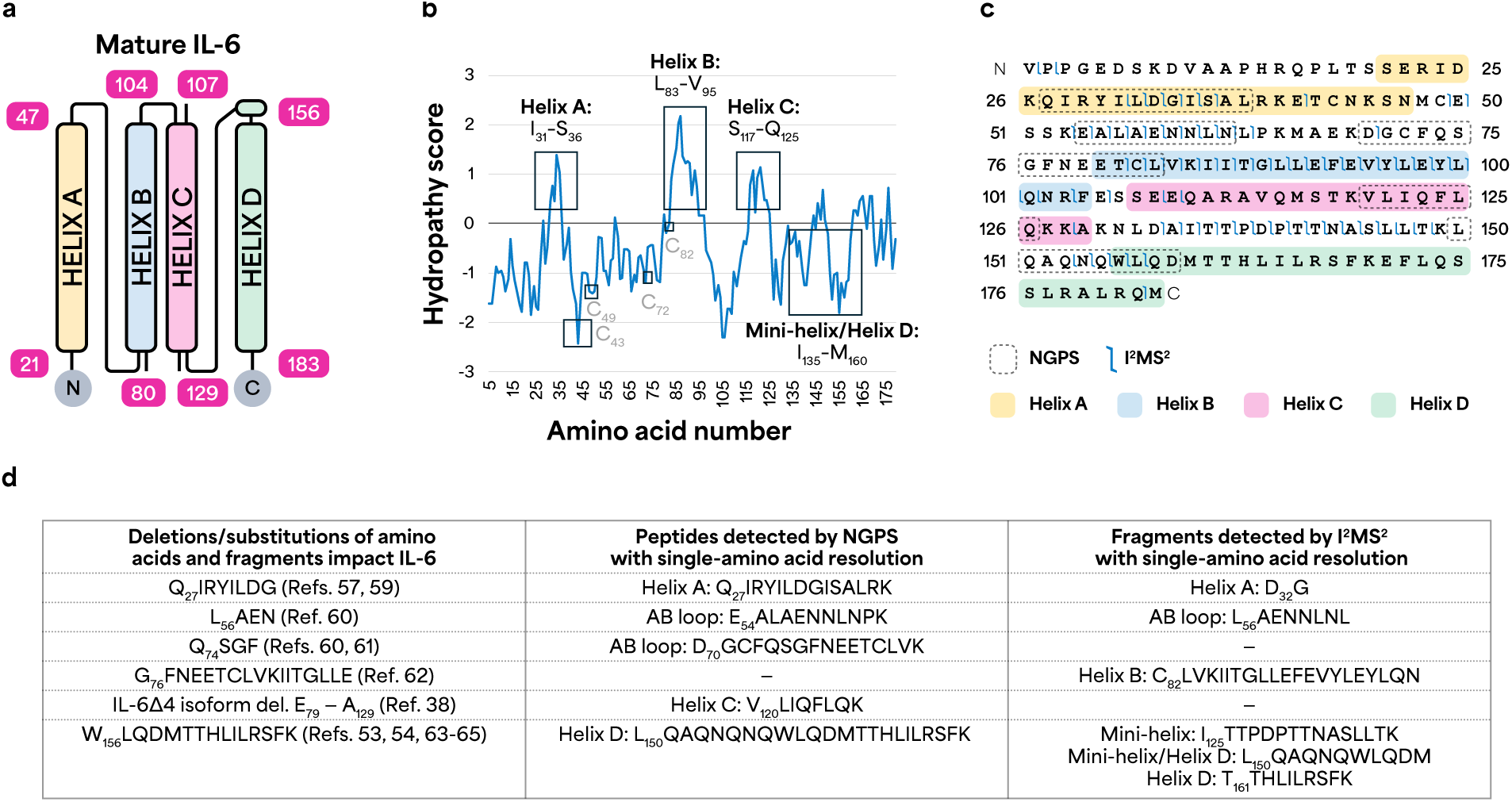
NGPS and I^2^MS/I^2^MS^2^ cover hydrophobic and hydrophilic regions of IL-6 important for IL-6 interactions. (**a**) Mature human IL-6 schematic. Numbering is based on a reported three-dimensional structure of human IL-6. (**b**) Kyte-Doolittle hydropathy plot reports surface-exposed regions. Positive scores correspond to hydrophobic regions. Negative scores correspond to hydrophilic regions. Amino acid sequence position is on the x axis. Average hydropathy score is calculated for windows of 9 amino acids. (**c**) NGPS (Platinum) and I^2^MS provide broad coverage of IL-6, combining to resolve 52% of single amino acids. (**d**) NGPS (Platinum) and I^2^MS detect single amino acids and protein fragments (denoted with numbers in columns 2 and 3) reported to impact IL-6 interactions (denoted with numbers in column 1).

IL-6, a four-helix bundle cytokine that contains 183 amino acids after signal peptide cleavage [58–60], is a multi-functional protein that transmits cellular signaling via IL-6 receptor alpha (IL-6R) and beta (gp130) [61]. The molecular information for IL-6 binding and activity is enabled via adoption of a four-helix fold, with a mini-helix before Helix D [57]. Each alpha helix contains 20-30 amino acids, with long AB and CD loops that accommodate an up-up-down-down helical orientation (**Figure 3A**). Previous studies indicate that multiple segments within IL-6 topology are necessary for biological interactions [62]. Hence, broad coverage of IL-6 sequence is key for elucidating residues important for IL-6 interactions, which occur via hydrophobic and hydrophilic interactions spread across different domains.

To determine if NGPS and I^2^MS provide broad coverage of the helices and loop regions, we generated a Kyte-Doolittle hydropathy plot [63] (**Figure 3B**). Interestingly, NGPS and I2MS analyze broad regions of IL-6 that are hydrophobic (positive scores) and hydrophilic (negative scores) (**Figure 3B**). NGPS resolves single amino acids within peptide Q27IRYILDGISAL38 (**Figure 3C**), located within Helix A, which is one of the most hydrophobic regions in IL-6 (**Figure 3B**). Further, NGPS also detects V120LIQFLQK127 located within a hydrophobic portion of Helix C (**Figure 3B**).

The most hydrophobic portion of IL-6 encompasses C82LVKIITGLLEFEVYLEYLQNR103 within Helix B (**Figure 3B**), an internal fragment that I^2^MS detects (**Figure 3C**). While this fragment is nonpolar, other cysteine-containing segments lie within hydrophilic regions (**Figure 3B**). I^2^MS also discerns N131LDAIITTPDPTTNASLLTK150 located within an amphipathic region of the CD loop and the mini helix that precedes Helix D (**Figures 3A, C**). Both I^2^MS and NGPS cover amino acids within Helix D (**Figure 3C**), a C-terminal region that contains a stretch of amphipathic residues (**Figure 3B**). Overall, NGPS and I^2^MS sequence various regions of IL-6 with different amino acid compositions and properties, highlighting the complementarity of these orthogonal analytical methods (**Figure 3C**).

For sequence structure-function analysis, we performed a literature survey to determine which sequenced regions are relevant to IL-6 interactions. Removal of amino acids within the Q27IRYILD33 segment of Helix A reduces or abolishes IL-6 activity [62, 64] (**Figure 3D, Row 1**). Within the AB loop, E54ALAEAENNLNL65 contains residues important for IL-6 binding to IL-6R [65] (**Figure 3D, Row 2**). Similarly, D70GCFQSGFNE79 harbors residues such as phenylalanine (F), both of which are relevant for IL-6 binding interactions (**Figure 3D, Row 3**)[65–67]. We demonstrate that NGPS resolves both F residues within D70GCFQSGFNE79 (**Figure 1E**, **Figure 3C**). NGPS also detects peptide V120LIQFLQK127 within Helix C (**Figure 1E**, **Figure 3C**). Notably, a segment including V120LIQFLQK127 is truncated in an IL-6 splice variant with altered signaling (**Figure 3D, Row 4**) [33]. The mini-helix and amphipathic D helix contain a stretch of leucine and polar Q/N residues that influence IL-6 binding [55, 56, 68–70]. Therefore, both NGPS and I^2^MS detect regions of IL-6 relevant to its biological function and interactions.

Limitations of this study include the use of recombinant IL-6 and, as a proof-of-concept study, its small sample size. Future studies will aim to expand the scope of analysis for combining NGPS and I^2^MS^2^ across a broader range of target proteins and proteoforms.

## Conclusion

Taken together, our data demonstrate complementarity of single-molecule protein sequencing (NGPS on Platinum) and I^2^MS to cover key regions of IL-6. While NGPS provided single amino acid resolution for fragments like V120LIQFLQK127, which are essential for IL-6 receptor binding, I^2^MS enabled detection of larger proteoforms, providing critical information on PTMs, such as glycosylation, that affect IL-6’s stability and bioactivity. When combined, NGPS and I^2^MS cover many key regions of IL-6, achieving 52% combined sequence coverage of single amino acids. This sets up improvements in technology and informs future studies of how polymorphisms or mutations affect PTMs on endogenous IL-6. These complementary datasets underscore the potential of combining single-molecule protein sequencing and mass spectrometry to obtain a more comprehensive picture of IL-6 structural and functional diversity, which is vital for understanding its therapeutic potential.

## Author Contributions

Conception and project design: KAS, MAC, NLK

Data acquisition: KAS, TDF, AL, TS, EF, AS

Data analysis and interpretation of results: KAS, TDF, MW, HH, TS, EF, AS, MAC, NLK

Original draft preparation: KAS and MAC

All authors reviewed the results and approved the final version of the manuscript.

## Acknowledgments

We thank researchers at Quantum-Si, particularly Brian Reed and Meredith Carpenter, and members of the Northwestern Proteomics Center of Excellence (PCE) for technical assistance. Research reported in this publication was supported by the National Institute of General Medical Sciences of the National Institutes of Health under Award Number P41GM108569. The content is solely the responsibility of the authors and does not necessarily represent the official views of the National Institutes of Health.

## Supplementary Information

**Supplementary Figure 1.**
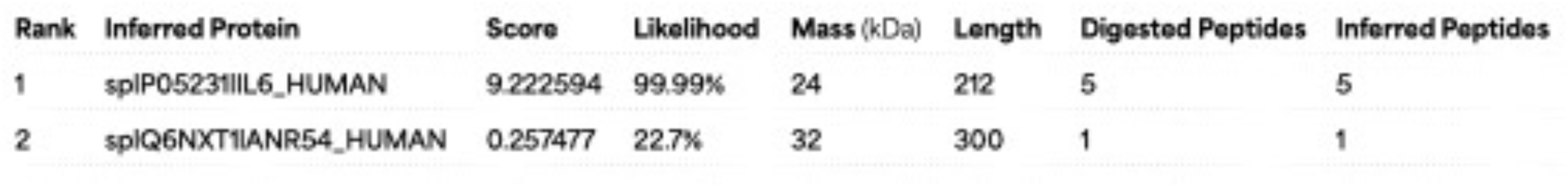
Protein inference v2.5.2 aligns observed kinetic signatures from sequencing data of rhIL-6 to a database of predicted kinetic signatures. Protein Score is an estimate of the likelihood of protein identity. While ankyrin repeat domain-containing protein 54 (UniProt ID: Q6NXT1) contains peptide QIIHMLREYLERLGQHEQRERLDDLCTRLQMTSTK, which is similar in sequence to QIRYILDGISALRK in rhIL-6, Platinum unambiguously identifies rhIL-6.

**Supplementary Table 1.** Intact Proteoforms of rhIL-6 detected with I. ^2^**MS.** 13 proteoforms of rhIL-6 were detected by I^2^MS and abundant species were targeted for tandem-MS with I^2^MS^2^. Based on the intact masses, putative glycan compositions were proposed and supported with I^2^MS^2^ data when available.

| PFR | Theoretical Monoisotopic Mass (Da) | Observed Monoisotopic Mass (Da) | Mass Error (ppm) | Half Cystine Sites | Putative O-Glycan Composition | Glycan Chemical Formula (unbound) | Monoisotopic Glycan Mass (Da) | I<sup>2</sup>MS<sup>2</sup>? |
| --- | --- | --- | --- | --- | --- | --- | --- | --- |
| 1 | 20795.58 | 20795.55 | -1.55 | C72, C78, C101, C111 | N/A | N/A | N/A | Yes |
| 2 | 20998.66 | 20998.64 | -1.03 | C72, C78, C101, C111 | HexNAc(1) | C8H13NO5 | +221.08994 | No |
| 3 | 21160.71 | 21160.69 | -1.16 | C72, C78, C101, C111 | HexNAc(1) Hex(1) | C14H23NO10 | +383.14276 | No |
| 4 | 21289.76 | 21289.72 | -1.74 | C72, C78, C101, C111 | HexNAc(1) NeuAc(1) | C19H30N2O13 | +512.18535 | Yes |
| 5 | 21452.17 | 21451.79 | -17.71 | C72, C78, C101, C111 | HexNAc(1) Hex(1) NeuAc(1) | C25H40N2O18 | +674.23818 | Yes |
| 6 | 21646.87 | 21646.84 | -1.52 | C72, C78, C101, C111 | HexNAc(1) Hex(4) | C32H53NO25 | +869.30123 | No |
| 7 | 21654.89 | 21654.86 | -1.35 | C72, C78, C101, C111 | HexNAc(2) Hex(1) NeuAc(1) | C33H53N3O23 | +877.31755 | No |
| 8 | 21742.91 | 21742.90 | -0.25 | C72, C78, C101, C111 | HexNAc(1) Hex(1) NeuAc(2) | C36H57N3O26 | +965.33359 | Yes |
| 9 | 21785.95 | 21785.89 | -2.64 | C72, C78, C101, C111 | HexNAc(4) Hex(1) | C38H62N4O25 | +992.38088 | Yes |
| 10 | 21858.99 | 21858.91 | -3.62 | C72, C78, C101, C111 | HexNAc(3) dHex(2) Hex(1) | C42H69N3O28 | +1081.41732 | No |
| 11 | 21905.98 | 21905.95 | -1.30 | C72, C78, C101, C111 | HexNAc(1) dHex(2) Hex(2) NeuAc(1) | C43H70N2O31 | +1128.40682 | Yes |
| 12 | 21947.01 | 21946.97 | -1.60 | C72, C78, C101, C111 | HexNAc(2) dHex(2) Hex(1) NeuAc(1) | C45H73N3O31 | +1169.43337 | Yes |
| 13 | 22109.06 | 22109.03 | -1.26 | C72, C78, C101, C111 | HexNAc(2) dHex(2) Hex(2) NeuAc(1) | C51H83N3O36 | +1331.48619 | Yes |

**Supplementary Figure 2.**
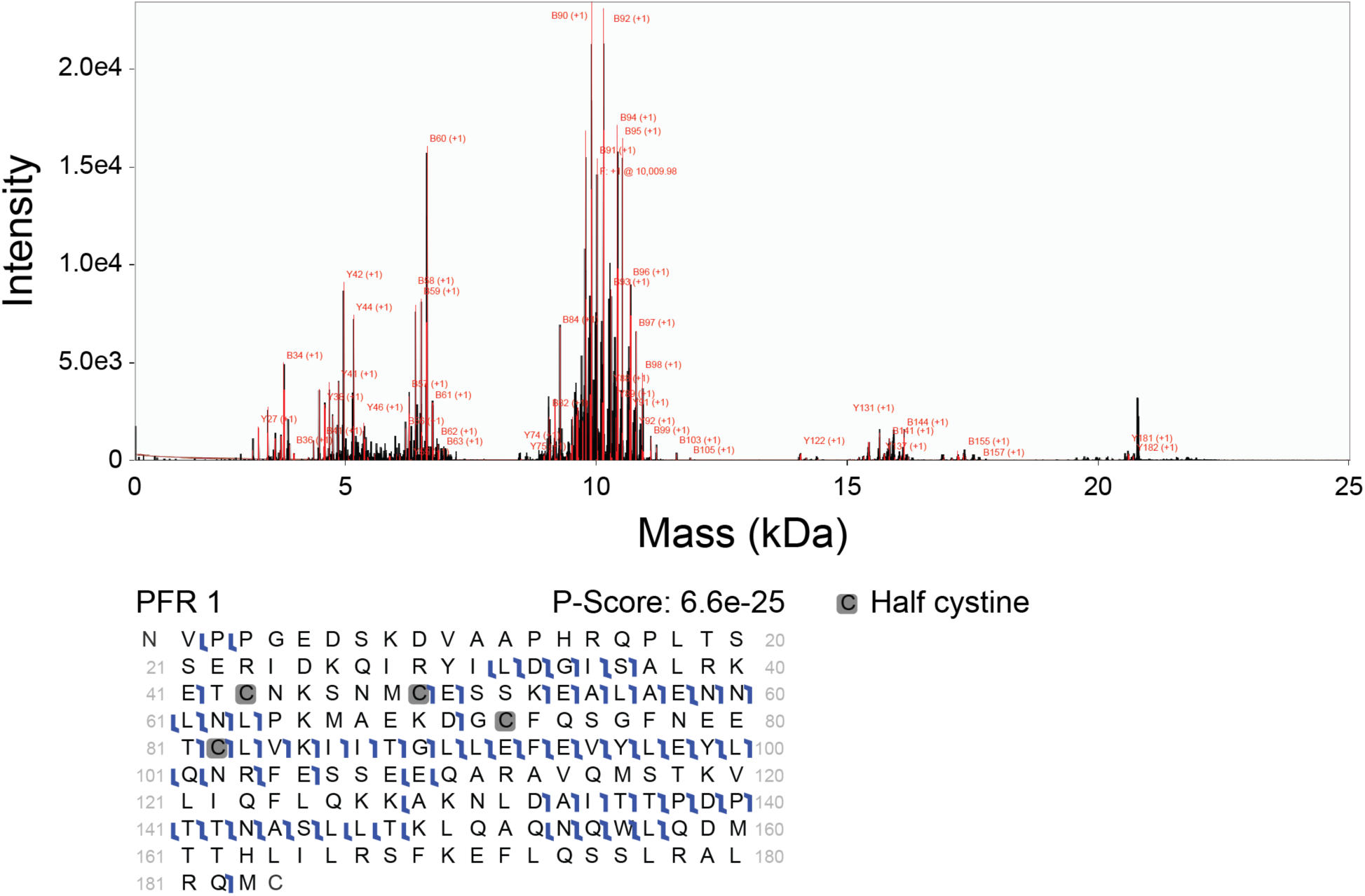
Fragmentation spectrum and graphical fragment map for rhIL-6 proteoform 1.

**Supplementary Figure 3.**
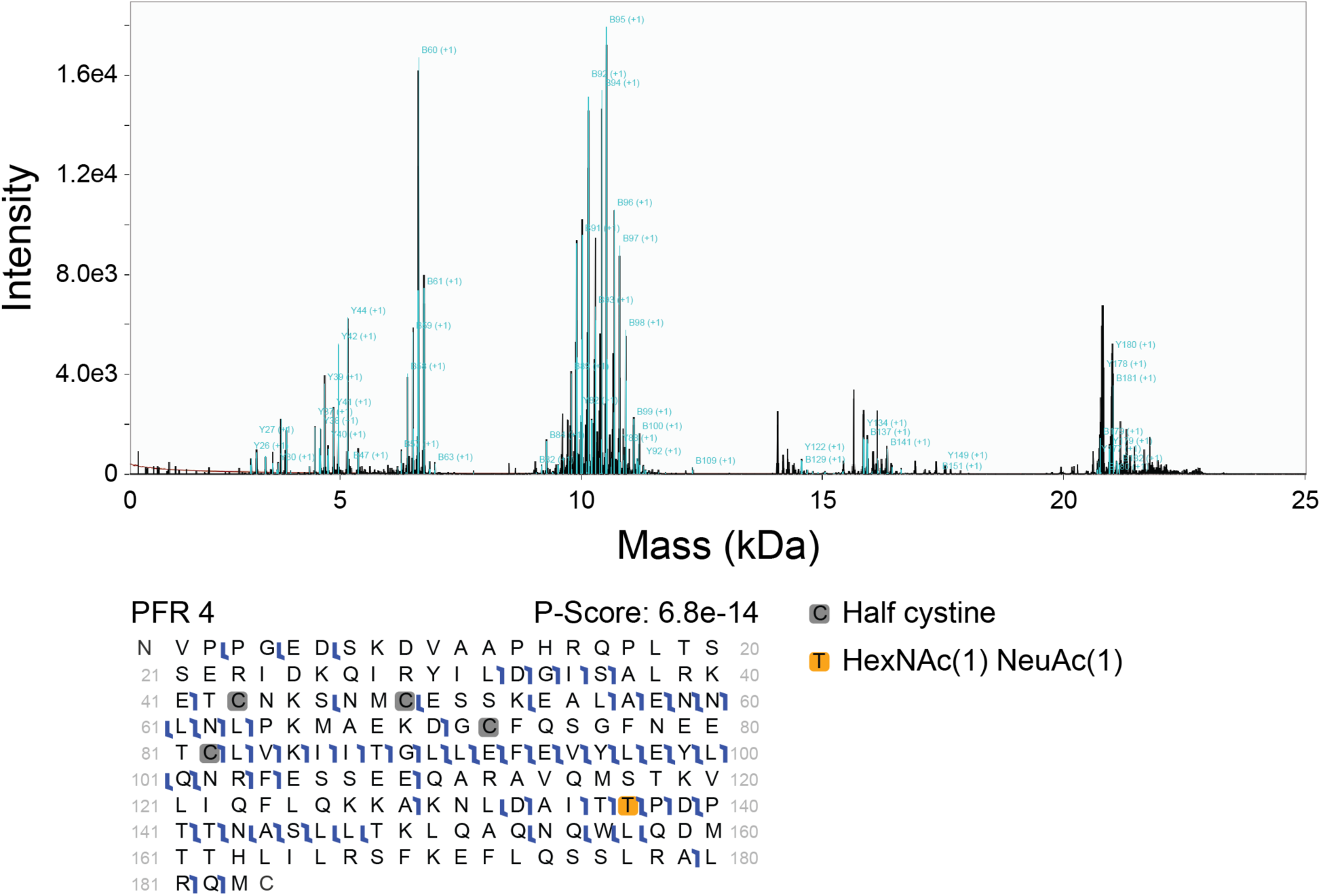
Fragmentation spectrum and graphical fragment map for rhIL-6 proteoform 4.

**Supplementary Figure 4.**
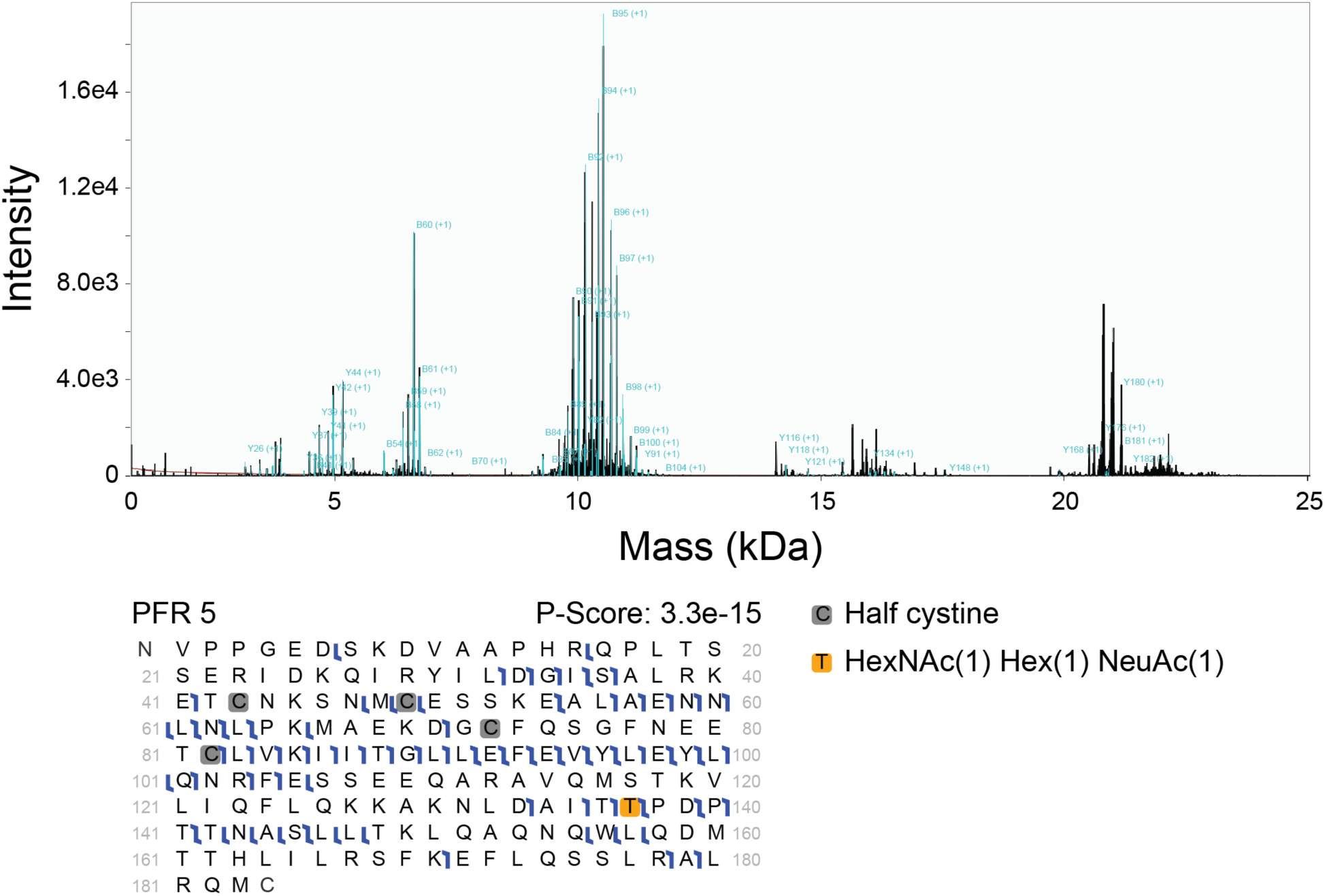
Fragmentation spectrum and graphical fragment map for rhIL-6 proteoform 5.

**Supplementary Figure 5.**
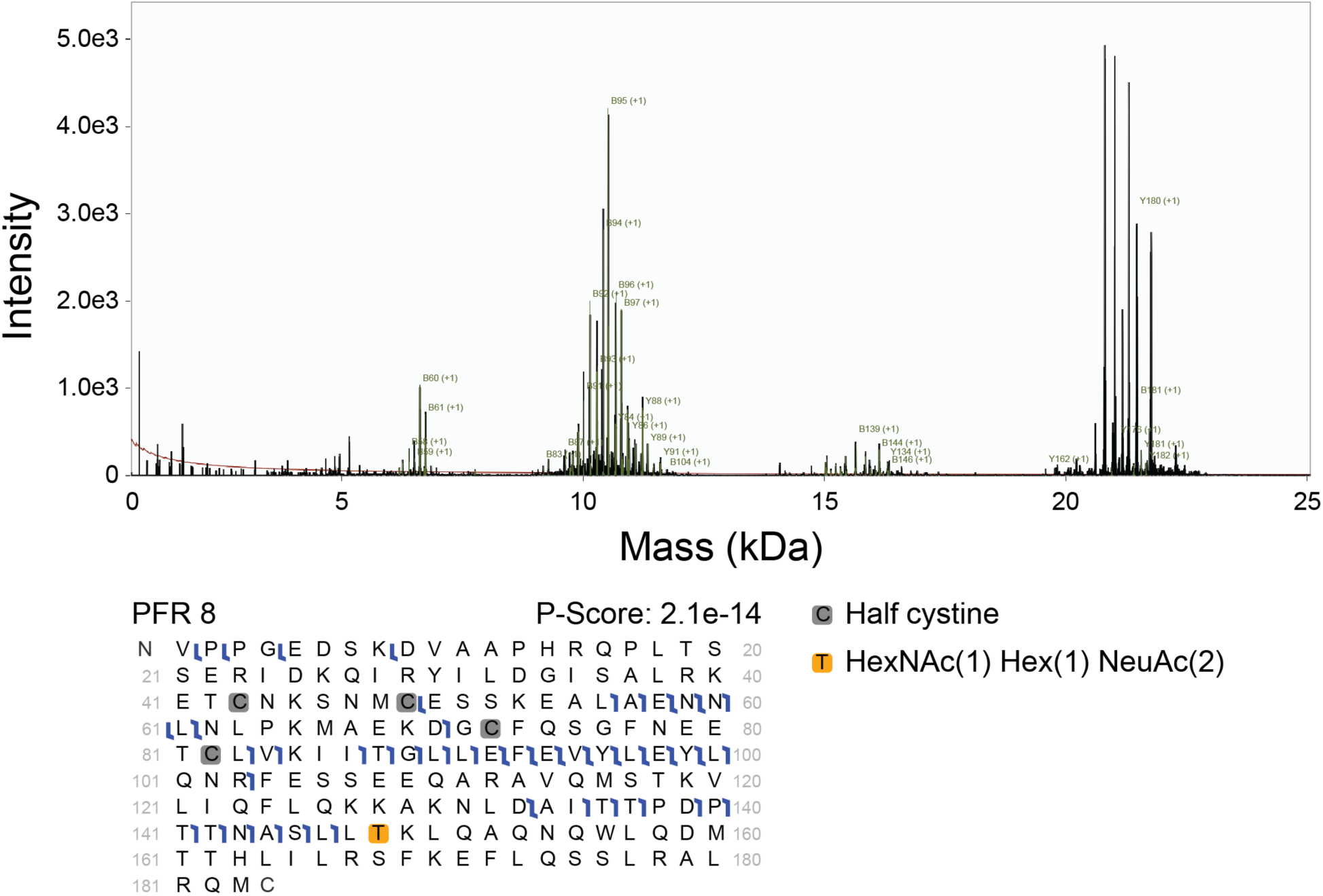
Fragmentation spectrum and graphical fragment map for rhIL-6 proteoform 8.

**Supplementary Figure 6.**
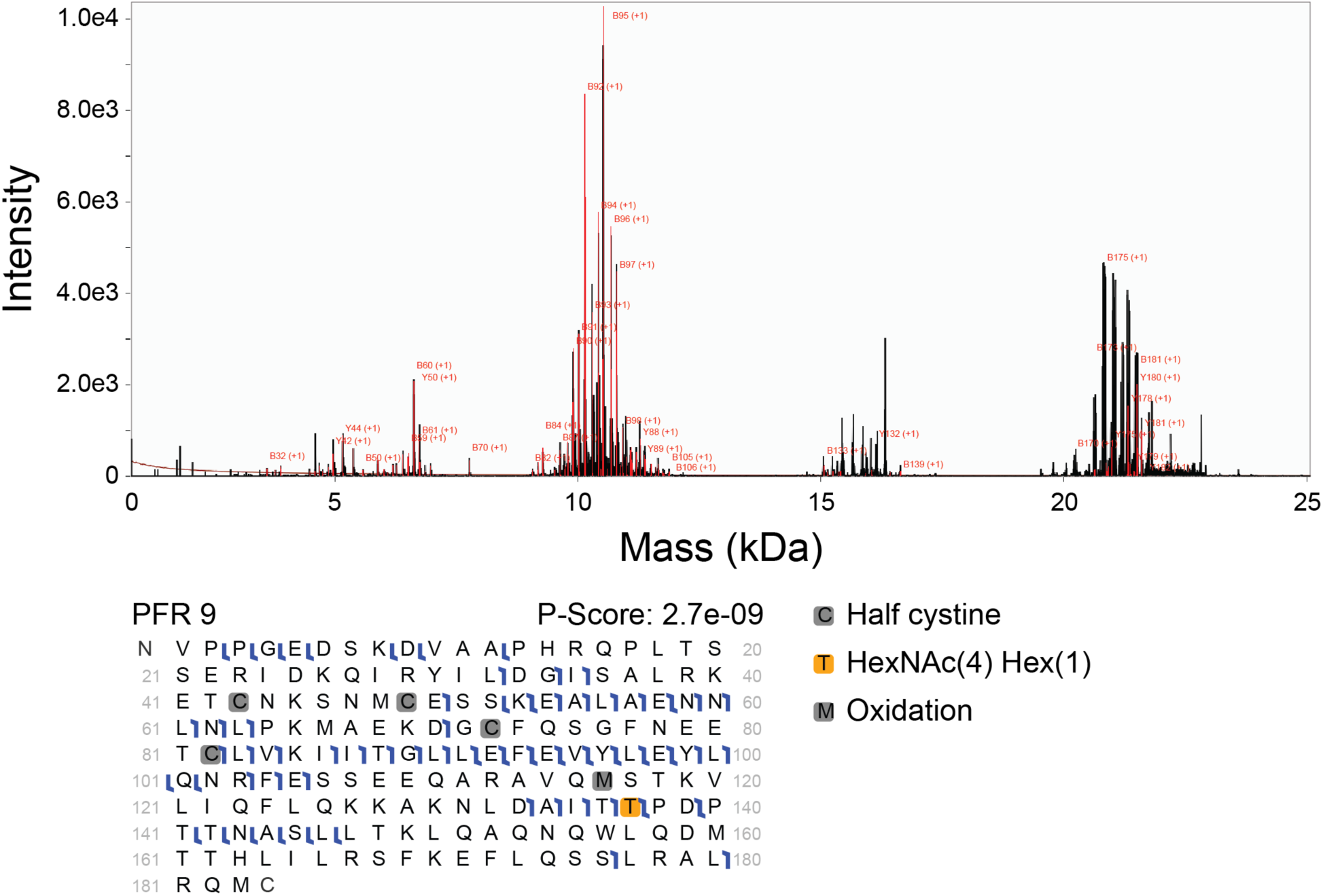
Fragmentation spectrum and graphical fragment map for rhIL-6 proteoform 9.

**Supplementary Figure 7.**
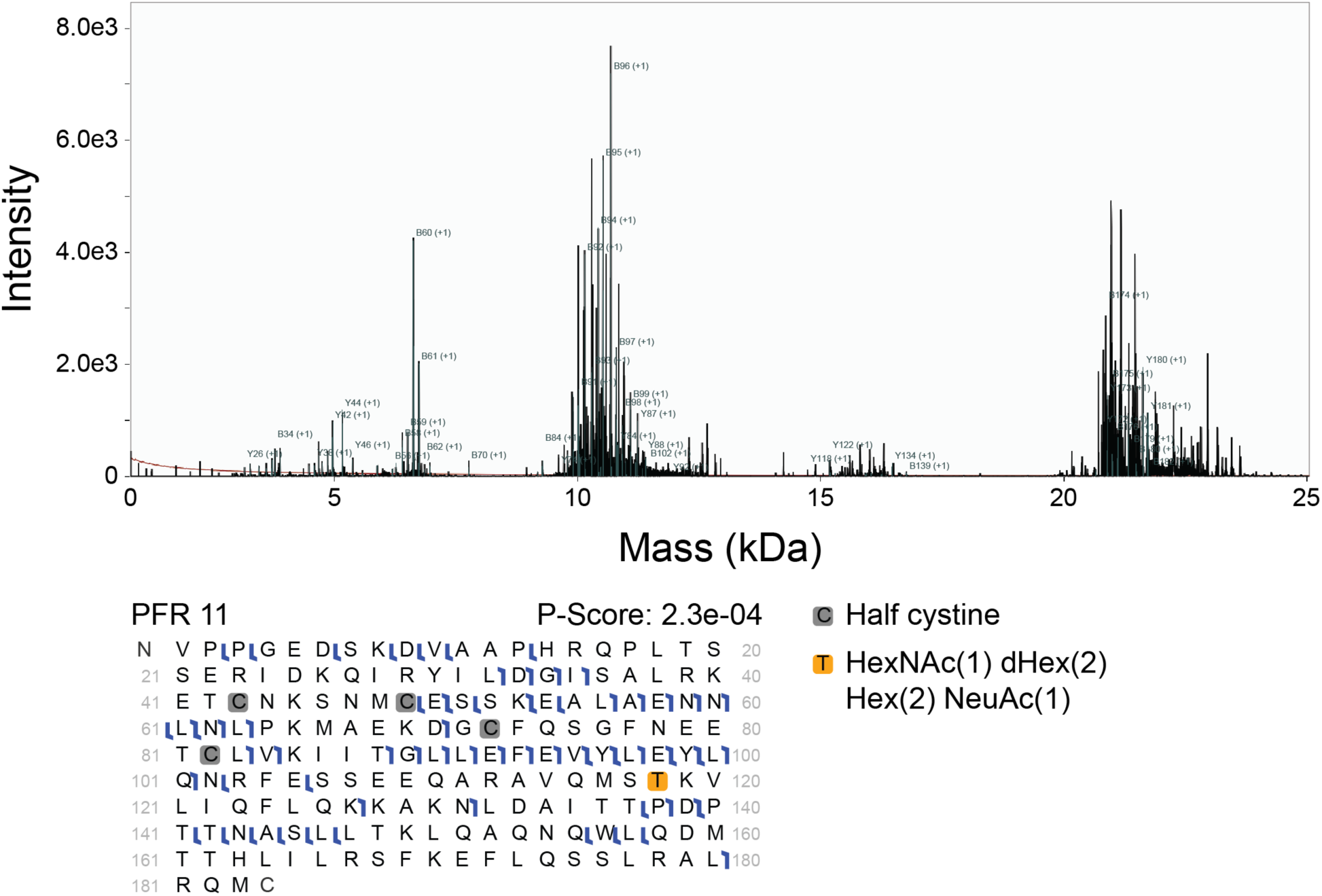
Fragmentation spectrum and graphical fragment map for rhIL-6 proteoform 11.

**Supplementary Figure 8.**
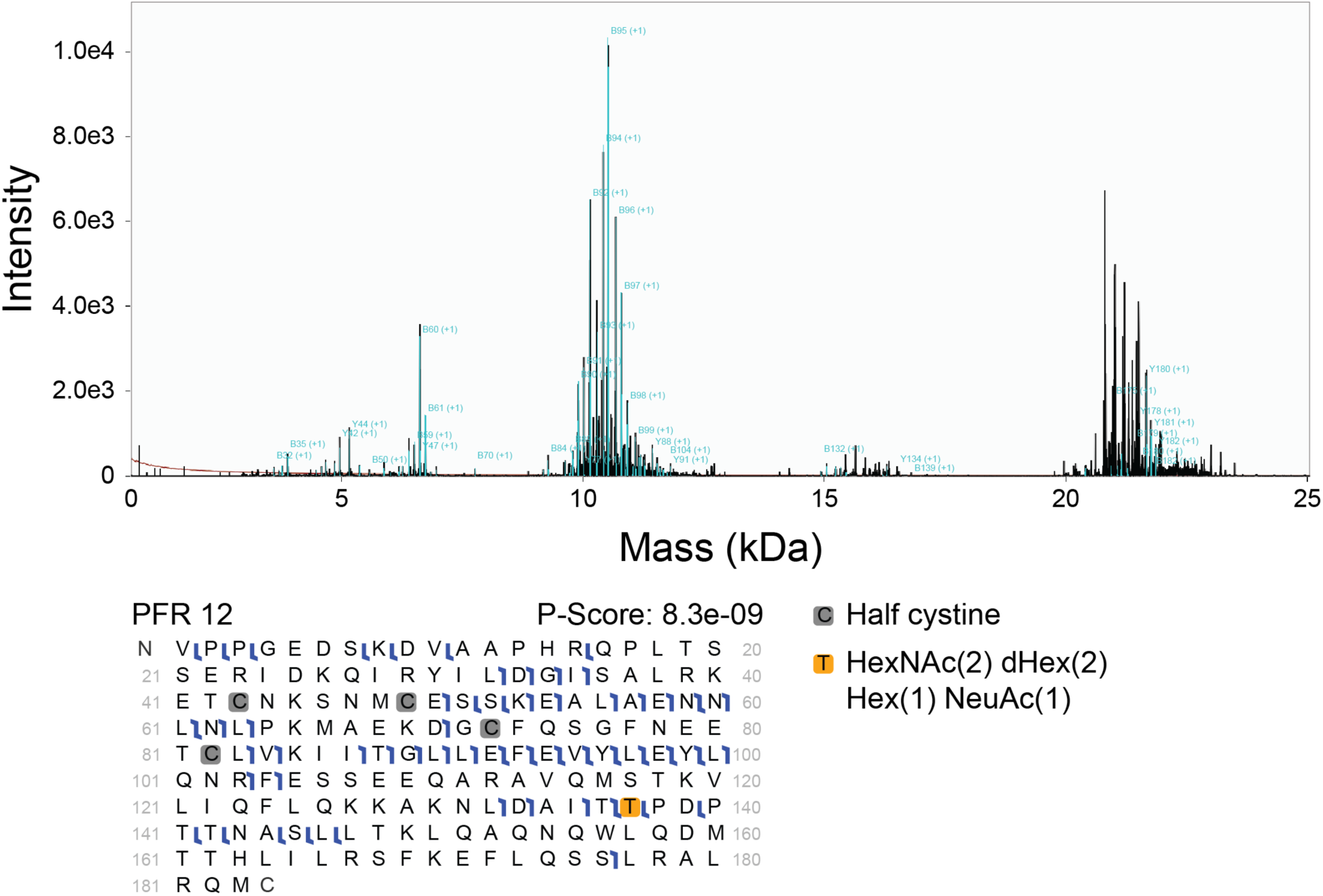
Fragmentation spectrum and graphical fragment map for rhIL-6 proteoform 12.

**Supplementary Figure 9.**
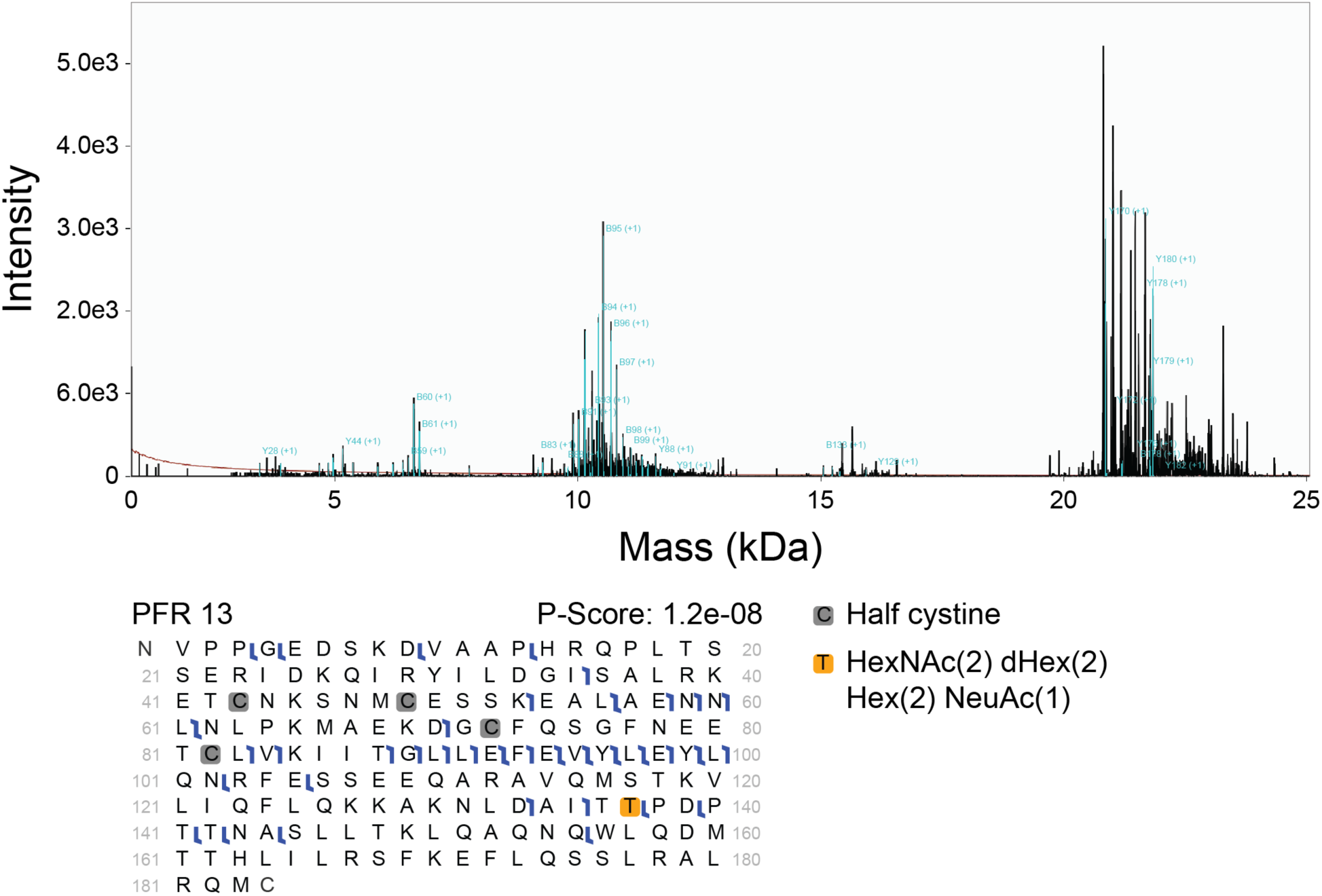
Fragmentation spectrum and graphical fragment map for rhIL-6 proteoform 13.

